# Myeloid mDIC Deficiency Remodels Adipose Macrophages to an Efferocytosis-Associated State and Improves Metabolic Homeostasis During Obesity

**DOI:** 10.64898/2026.11.17.767460

**Authors:** Fenfen Wang, Phu Minh Huynh, Alexandra Serrato, Qixiu Hou, Yang Yang, Xinxin Ma, Juli Bai, Yu A. An

## Abstract

Obesity-associated adipose inflammation contributes to metabolic dysfunction, yet the mechanisms linking mitochondrial metabolism to macrophage efferocytosis remain incompletely understood. The mitochondrial dicarboxylate carrier (mDIC), encoded by *SLC25A10*, regulates adipocyte function, but its role in macrophages remains unclear. Here, we investigated how myeloid mDIC deficiency affects mitochondrial function, macrophage heterogeneity, and metabolic homeostasis during diet-induced obesity. We observed reduced mDIC protein expression in subcutaneous white adipose tissue from individuals with obesity, whereas palmitate increased mDIC expression in mouse and human macrophages. Myeloid mDIC deletion preserved mitochondrial respiration under palmitate stress. Single-cell RNA sequencing of adipose stromal vascular cells from high-fat diet (HFD)-fed mice revealed an expanded efferocytosis-associated macrophage population and reduced proportions of inflammatory and lipid-associated macrophages. Consistently, mDIC-deficient macrophages exhibited increased *Mertk* expression and greater association with apoptotic cells. Pharmacological inhibition of mitochondrial ATP synthase or electron transport markedly suppressed apoptotic cell uptake, including the quenching-resistant fraction, without substantially affecting macrophage viability. *In vivo*, myeloid mDIC deletion attenuated HFD-induced weight gain, improved insulin sensitivity, and increased CD11c⁻CD206⁺ adipose macrophage abundance, accompanied by elevated expression of alternative activation-associated markers. Together, these findings identify myeloid mDIC as an important regulator of macrophage mitochondrial function and adipose immune remodeling during obesity, suggesting a novel mechanism by which mitochondrial bioenergetics contribute to macrophage apoptotic-cell handling and metabolic homeostasis.

## Introduction

Obesity is a major risk factor for type 2 diabetes and cardiovascular disease and is associated with chronic low-grade inflammation and metabolic dysfunction^1–7^. Adipose tissue plays a central role in obesity-associated inflammation, where excessive lipid accumulation, adipocyte hypertrophy, and cell death promote immune cell recruitment and tissue remodeling^1,8–10^. Among adipose immune cells, adipose tissue macrophages (ATMs) are key regulators of tissue homeostasis and metabolic inflammation. Recent single-cell transcriptomic studies have revealed substantial ATM heterogeneity, identifying distinct macrophage populations involved in lipid handling, inflammatory responses, and tissue remodeling^1,11–16^. Notably, lipid-associated macrophages (LAMs) exhibit specialized lipid metabolic programs and contribute to adipose tissue adaptation during obesity^11^. These findings highlight the functional diversity of ATMs and suggest that changes in macrophage composition and activity may influence obesity-associated metabolic dysfunction.

Efferocytosis, the recognition and clearance of apoptotic cells by phagocytes, is essential for maintaining tissue homeostasis and promoting inflammatory resolution^17–23^. During obesity, increased adipocyte death creates a sustained demand for macrophage-mediated clearance of dying cells^8,13,17^. Efficient efferocytosis prevents secondary necrosis and limits the release of pro-inflammatory cellular contents, whereas defective clearance can contribute to persistent tissue inflammation^17–19,24^. This process involves multiple coordinated steps, including apoptotic-cell recognition, binding, internalization, phagosome maturation, and intracellular degradation^18,25,26^. Several receptors participate in apoptotic-cell clearance, including Mer tyrosine kinase (MerTK), which plays an important role in macrophage efferocytosis^18,27–29^. Recent evidence further links macrophage efferocytosis to the resolution of adipose inflammation. For example, deletion of the miR-130b/301b cluster enhances macrophage efferocytosis and is associated with reduced adipose inflammatory gene expression during high-fat diet (HFD) feeding^30^. However, the mechanisms regulating the development and function of efferocytosis-associated ATMs during obesity remain poorly understood.

Growing evidence indicates that mitochondrial metabolism plays an important role in macrophage efferocytosis and inflammatory responses^31–33^. Mitochondrial fission facilitates continued apoptotic-cell clearance by regulating intracellular calcium signaling and phagosome formation^34^. Importantly, engulfment of apoptotic cells has been shown to increase macrophage oxygen consumption rate (OCR), linking efferocytosis to mitochondrial respiratory activity^35^. Consistently, a recent study demonstrated that efferocytosis increased basal and ATP-linked OCR, while deletion of the miR-130b/301b cluster further enhanced mitochondrial respiration and efferocytic capacity^30^. In addition, mitochondrial fatty acid oxidation and electron transport support efferocytosis-induced IL-10 production and tissue repair^35^. Efferocytosis also induces metabolic reprogramming involving solute carrier-mediated glucose uptake and glycolysis, which contribute to apoptotic-cell handling and anti-inflammatory responses^36^. More recently, mitochondrial ATP production was implicated in the regulation of macrophage efferocytosis^37^. Together, these findings highlight the importance of metabolic adaptation in macrophage apoptotic-cell clearance, although the contribution of mitochondrial respiration may vary across experimental contexts and stages of efferocytosis. However, whether mitochondrial metabolite transport regulates macrophage respiratory function and efferocytosis during obesity remains largely unexplored.

The mitochondrial dicarboxylate carrier (mDIC), encoded by *SLC25A10*, is an inner mitochondrial membrane transporter that mediates the exchange of dicarboxylates, including malate and succinate, across the mitochondrial membrane^38,39^. Through mitochondrial metabolite transport, mDIC participates in metabolic processes associated with lipid synthesis and cellular metabolic homeostasis^38^. Previous studies by our group demonstrated that mDIC regulates adipocyte lipid metabolism, with its deficiency promoting adipocyte lipolysis and hepatic lipotoxicity^40^. Moreover, reduced adipose mDIC expression in humans has been associated with insulin resistance and metabolic dysfunction^40^. Although these studies establish an important role for mDIC in adipocyte metabolism, its function in macrophages remains uncharacterized. Given the metabolic plasticity of ATMs and the involvement of mitochondrial function in apoptotic-cell clearance, mDIC may represent an important link between mitochondrial metabolite transport and macrophage responses during obesity.

In the present study, we investigated the role of myeloid mDIC in macrophage mitochondrial function, adipose immune remodeling, and metabolic homeostasis during diet-induced obesity. We examined mDIC expression in human adipose tissue and palmitate-treated mouse and human macrophages and generated myeloid-specific mDIC knockout (mDIC MKO) mice to investigate its physiological function. Using mitochondrial respiration assays, single-cell RNA sequencing, apoptotic cell association assays, and HFD-induced obesity models, we evaluated how mDIC deficiency affects macrophage metabolism and the immune composition of adipose tissue. We further examined the contribution of mitochondrial activity to apoptotic-cell uptake using pharmacological inhibitors of ATP synthase and electron transport. Our findings identify myeloid mDIC as a critical regulator of macrophage mitochondrial responses and adipose immune remodeling, providing novel insights into the potential relationship between mitochondrial metabolite transport, efferocytosis-associated macrophage states, and metabolic homeostasis during obesity.

## Results

### 1. Palmitate induces mDIC expression in mouse and human macrophages despite reduced adipose mDIC expression in obesity

Our previous study demonstrated that the mitochondrial dicarboxylate carrier (mDIC/SLC25A10) regulates adipocyte lipolysis and systemic metabolic homeostasis^40^. Notably, reduced mDIC expression in human white adipose tissue (WAT) was associated with insulin resistance and metabolic dysfunction. Consistent with these findings, we observed significantly lower mDIC protein levels in subcutaneous WAT (SWAT) from individuals with obesity compared with lean individuals (P = 0.0263; Supplementary Fig. S1A).

Although the metabolic function of mDIC in adipocytes has been established, its regulation in adipose tissue macrophages (ATMs) remains largely unexplored. To investigate whether metabolic stress affects macrophage mDIC expression, we treated mouse bone marrow-derived macrophages (BMDMs) and human THP-1-derived macrophages with palmitate (PA) to mimic high-fat diet (HFD)-associated fatty acid exposure. Interestingly, PA treatment significantly increased both *Slc25a10* mRNA (P = 0.0272) and mDIC protein expression (P = 0.0224) in BMDMs (Fig. 1A). Similarly, PA increased *mDIC* mRNA (P = 0.0056) and mDIC protein levels (P = 0.0012) in THP-1-derived macrophages (Fig. 1B). Together, these findings reveal differential regulation of mDIC in whole adipose tissue during obesity and macrophages following PA exposure, suggesting that mDIC may have distinct roles in adipocytes and macrophages under metabolic stress.

**Figure 1.**
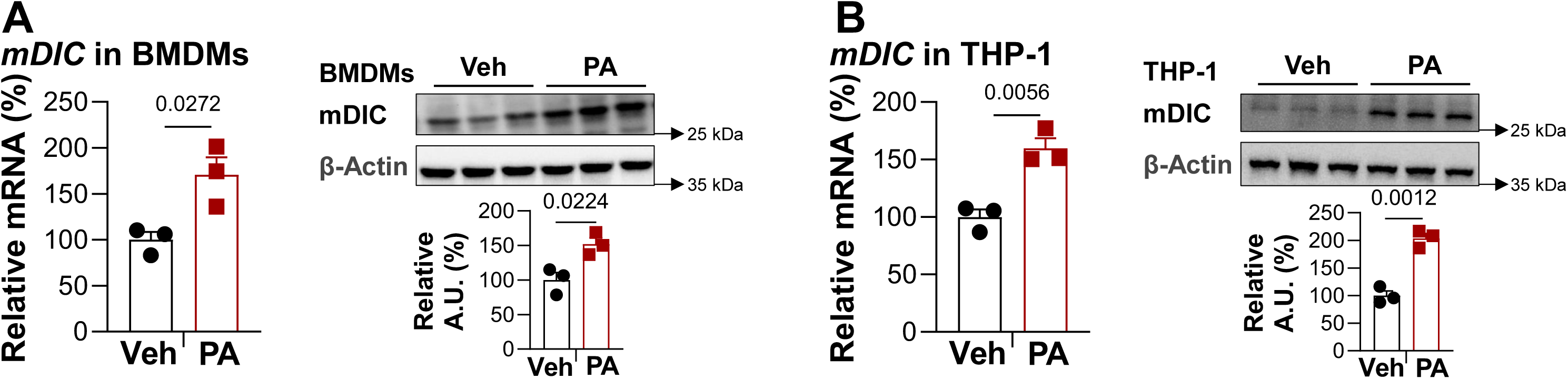
Palmitate increases mDIC expression in mouse and human macrophages. (A) Relative *mDIC* mRNA expression, representative mDIC and β-actin immunoblots, and densitometric quantification in mouse bone marrow-derived macrophages (BMDMs) treated with vehicle (Veh) or palmitate (PA; 100 μM, 24 h). (B) Relative *mDIC* mRNA expression, immunoblots, and quantification in human THP-1-derived macrophages treated with Veh or PA (100 μM, 24 h). Protein abundance was normalized to β-actin. Data are mean ± SEM (n = 3 per group). Two-tailed unpaired Student’s t-tests for A and B. Exact P values are shown.

### 2. Myeloid mDIC deletion enhances mitochondrial respiratory capacity under palmitate stress

To investigate the role of mDIC in macrophage mitochondrial function, we generated myeloid-specific *mDIC* knockout (mDIC MKO) mice by crossing *mDIC*-floxed mice with LysM-Cre mice^41^ (Fig. 2A). Successful deletion of mDIC was confirmed in bone marrow-derived macrophages (BMDMs), as demonstrated by significantly reduced *mDIC* mRNA (P < 0.0001; Fig. 2B) and mDIC protein expression (P = 0.0024; Fig. 2C).

**Figure 2.**
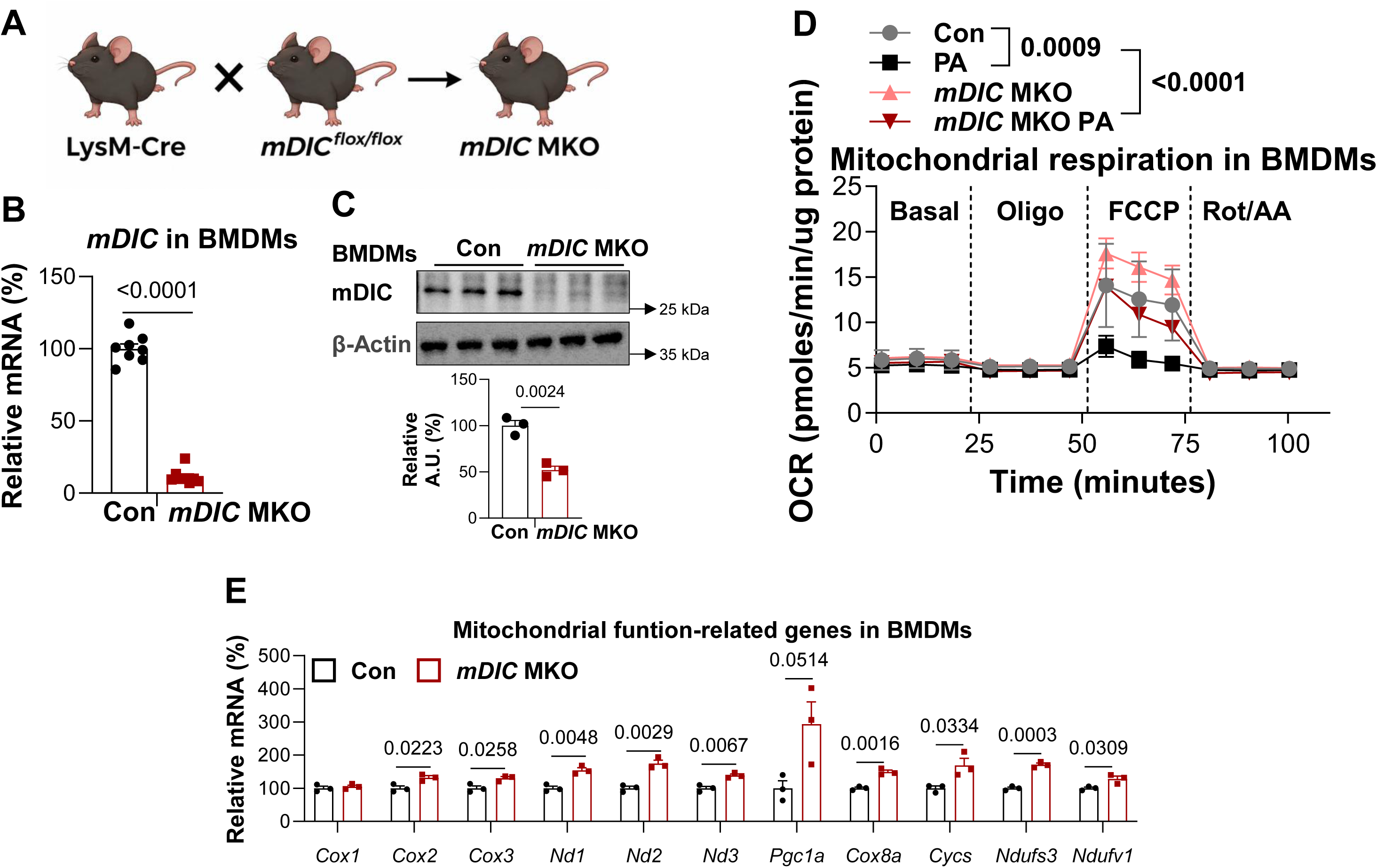
Myeloid mDIC deletion alters mitochondrial respiration and mitochondrial function-related gene expression. (A) Schematic showing generation of myeloid-specific *mDIC* knockout (*mDIC* MKO) mice by crossing LysM-Cre and *Slc25a10*-floxed mice. (B) Relative *mDIC* mRNA expression in control (Con) and *mDIC* MKO BMDMs (n = 8 per group). (C) Representative mDIC and β-actin immunoblots and densitometric quantification (n = 3 per group). (D) Seahorse oxygen consumption rate (OCR) profiles of Con and *mDIC* MKO BMDMs with or without PA treatment (100 μM, 24 h; n ≥ 4 per group). Oligomycin, FCCP, and rotenone/antimycin A (Rot/AA) were sequentially injected; OCR was normalized to total protein. (E) Relative mRNA expression of mitochondrial function-related genes (*Cox1*, *Cox2*, *Cox3*, *Nd1*, *Nd2*, *Nd3*, *Pgc1a*, *Cox8a*, *Cyc5*, *Ndufs3*, and *Ndufv1*) in in BMDMs derived from Con and *mDIC* MKO mice fed an HFD for 10 weeks (n = 3 per group). Data are mean ± SEM. Two-tailed unpaired Student’s t-tests for B, C, and E; two-way ANOVA for D. Exact P values are shown.

We next assessed mitochondrial respiration in control and mDIC MKO BMDMs following palmitate (PA) treatment using Seahorse extracellular flux analysis. Although basal oxygen consumption rates (OCRs) were comparable among groups, PA markedly suppressed FCCP-stimulated respiration in control BMDMs. In contrast, *mDIC* MKO BMDMs maintained higher FCCP-stimulated OCR following PA exposure (Fig. 2D), suggesting that mDIC deficiency preserves mitochondrial respiratory capacity under metabolic stress. Consistent with this observation, *mDIC* MKO BMDMs exhibited increased expression trends for multiple mitochondrial respiratory chain-related genes, including *Cox2*, *Nd1*, *Nd2*, *Nd3*, *Cox8a*, *Ndufs3*, and *Ndufv1*, as well as *Pgc1a*, a regulator of mitochondrial biogenesis (Fig. 2E). As a complementary inflammatory challenge, lipopolysaccharide (LPS) substantially reduced FCCP-stimulated respiration in BMDMs, including mDIC-deficient cultures (Supplementary Fig. S1B), suggesting that the respiratory phenotype is dependent on the metabolic or inflammatory stimulus. Together, these findings suggest that mDIC deficiency alters mitochondrial respiratory function and is associated with increased expression of mitochondrial metabolism-related genes in macrophages.

### 3. Myeloid mDIC deficiency remodels adipose tissue macrophage heterogeneity during early HFD feeding

To investigate how myeloid mDIC deficiency affects adipose immune cell populations during obesity, we performed single-cell RNA sequencing (scRNA-seq) of the stromal vascular fraction (SVF) isolated from subcutaneous white adipose tissue (SWAT) of control and *mDIC* MKO mice after 5 weeks of high-fat diet (HFD) feeding (Fig. 3A). Unsupervised clustering identified distinct cell populations, including B cells, T cells, CD4⁺ T cells, monocytes, macrophages, cycling cells, endothelial cells, fibroblasts, and mural cells (Fig. 3B). These populations were annotated based on established lineage markers, including *Ms4a1* and *Bank1* for B cells, *Cd247* and *Cd4* for T-cell populations, *Adgre1* for macrophages, *Lyz2* for myeloid cells, and *Mki67* for cycling cells (Supplementary Fig. S2A).

**Figure 3.**
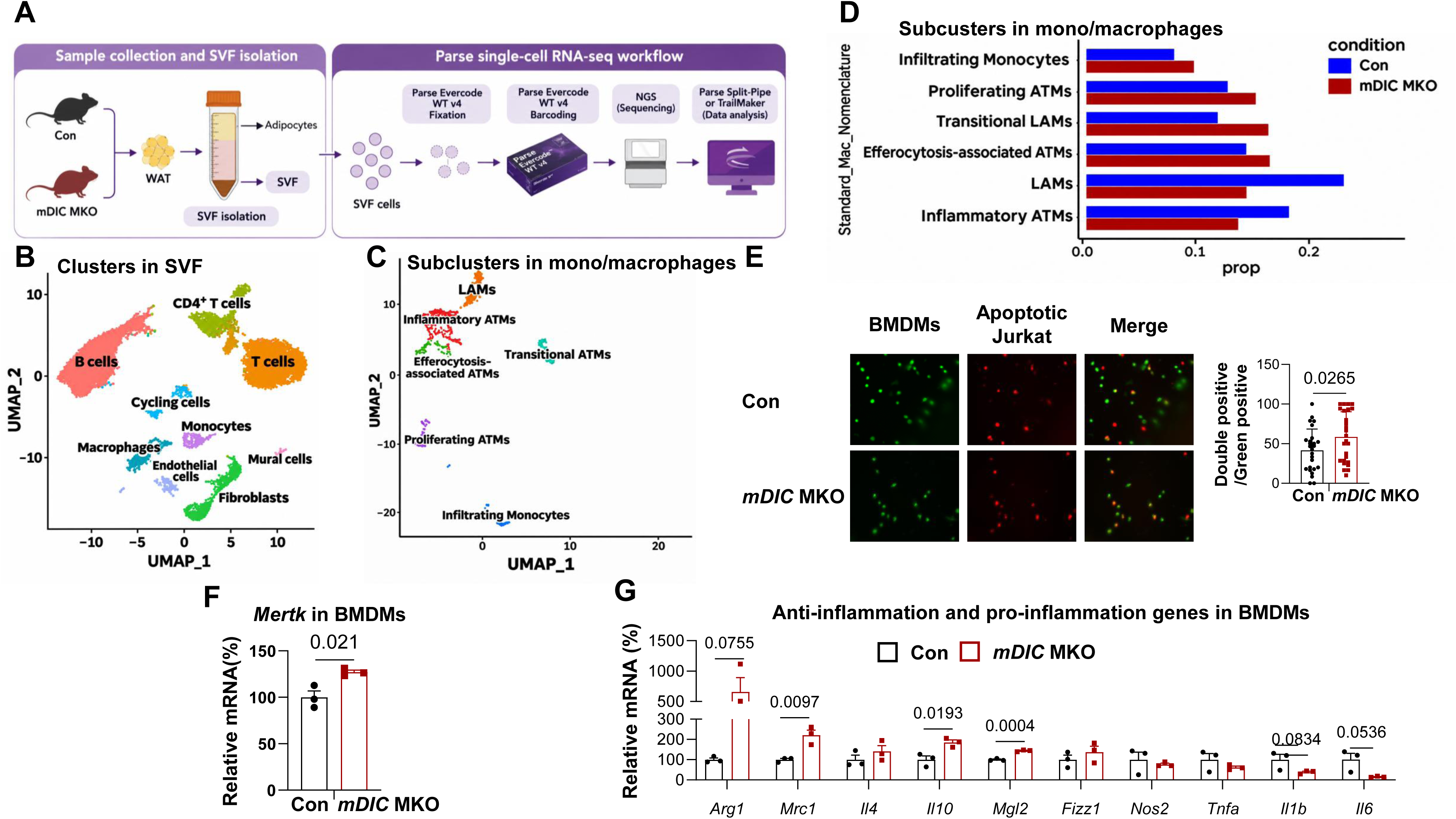
Myeloid mDIC deficiency remodels adipose macrophage heterogeneity and alters apoptotic-cell association and macrophage gene expression. (A) Schematic of stromal vascular fraction (SVF) isolation from SWAT of Con and *mDIC* MKO mice after 5 weeks of high-fat diet (HFD) feeding, followed by Parse single-cell RNA sequencing (scRNA-seq). Approximately 22,000–25,000 SVF cells were analyzed per genotype. (B) Uniform manifold approximation and projection (UMAP) showing the major SVF cell populations. (C) UMAP showing six monocyte/macrophage subclusters: inflammatory ATMs, lipid-associated macrophages (LAMs), efferocytosis-associated ATMs, transitional LAMs, proliferating ATMs, and infiltrating monocytes Approximately 1000 mono/macrophages were analyzed per genotype. (D) Relative proportions of these six subclusters in Con and *mDIC* MKO mice. Con and *mDIC* MKO mice were fed a HFD for 10 weeks, after which bone marrow cells were isolated and differentiated into BMDMs (E-G). (E) Representative fluorescence images of Con and *mDIC* MKO BMDMs (green) incubated with PKH26-labeled apoptotic Jurkat cells (red) at a 1:1 ratio for 30 min, with quantification of double-positive fluorescence relative to macrophage-associated green fluorescence (8 BMDM samples per group; at least three images per sample). (F) Relative *Mertk* mRNA expression in Con and *mDIC* MKO BMDMs (n = 3 per group). (G) Relative mRNA expression of alternative activation/anti-inflammatory-associated genes (*Arg1*, *Mrc1*, *Il4*, *Il10*, *Mgl2*, and *Fizz1*) and pro-inflammatory genes (*Nos2*, *Tnfa*, *Il1b*, and *Il6*) in Con and *mDIC* MKO BMDMs (n = 3 per group). Data are mean ± SEM. Two-tailed unpaired Student’s t-tests for E–G; Exact P values are shown where indicated.

To further characterize macrophage heterogeneity, we performed subclustering of the monocyte/macrophage compartment and identified six transcriptionally distinct populations: inflammatory ATMs, lipid-associated macrophages (LAMs), efferocytosis-associated ATMs, transitional LAMs, proliferating ATMs, and infiltrating monocytes (Fig. 3C). Marker gene analysis revealed that LAMs expressed lipid metabolism-associated genes, including *Pparg*, *Apoe*, and *Pla2g7*, whereas transitional LAMs exhibited expression of *Pparg* and *Lpl*. Notably, the efferocytosis-associated ATM population was characterized by prominent expression of *Mrc1* and *Lpl*, consistent with a macrophage state associated with tissue homeostasis and lipid handling. Proliferating ATMs were distinguished by expression of cell-cycle genes *Mki67* and *Top2a*, while infiltrating monocytes expressed *Lyz2* and *Ctsl* (Supplementary Fig. S2B).

Comparison of macrophage subcluster proportions revealed an increased relative abundance of efferocytosis-associated ATMs in *mDIC* MKO mice, accompanied by reduced proportions of LAMs and inflammatory ATMs (Fig. 3D). The relative proportions of infiltrating monocytes, proliferating ATMs, and transitional LAMs were also increased in *mDIC* MKO mice. Together, these findings indicate that myeloid mDIC deficiency is associated with remodeling of adipose macrophage heterogeneity during early HFD feeding, including enrichment of an ATM population exhibiting an efferocytosis-associated transcriptional phenotype.

### 4. Myeloid *mDIC* deficiency promotes an efferocytosis-associated macrophage phenotype

To determine whether the altered adipose tissue macrophage composition was accompanied by changes in macrophage efferocytic activity, we incubated control and *mDIC* MKO bone marrow-derived macrophages (BMDMs) with fluorescently labeled apoptotic Jurkat cells. Fluorescence microscopy revealed greater apoptotic-cell-associated signal in *mDIC* MKO BMDMs compared with controls (Fig. 3E). Quantification of double-positive fluorescence relative to macrophage-associated green fluorescence showed an increasing trend in *mDIC* MKO BMDMs (Fig. 3E), suggesting enhanced interactions with apoptotic cells following mDIC deletion.

We next examined whether mDIC deficiency altered macrophage expression of efferocytosis-and inflammation-associated genes. RT-qPCR analysis revealed increased *Mertk* expression in *mDIC* MKO BMDMs compared with controls (Fig. 3F), consistent with the enrichment of efferocytosis-associated ATMs identified by scRNA-seq. In addition, *mDIC* MKO BMDMs exhibited increased expression of alternative activation-associated genes, particularly *Arg1* and *Mrc1*, along with upward trends in *Il10* and other related markers (Fig. 3G). Conversely, inflammatory markers, including *Nos2*, *Tnfa*, *Il1b*, and *Il6*, generally exhibited lower expression in *mDIC* MKO BMDMs. Together, these findings suggest that myeloid mDIC deficiency promotes a macrophage transcriptional phenotype associated with apoptotic-cell clearance and reduced inflammatory activation.

### 5. Mitochondrial inhibition markedly suppresses apoptotic-cell uptake-associated fluorescence in macrophages

Given that myeloid mDIC deletion altered mitochondrial respiration and was associated with an efferocytosis-related macrophage phenotype, we next investigated whether mitochondrial activity contributes to apoptotic-cell uptake. Previous studies demonstrated that mitochondrial fission facilitates continued apoptotic-cell clearance^34^, whereas mitochondrial fatty acid oxidation and electron transport support efferocytosis-induced anti-inflammatory responses^35^. More recently, mitochondrial ATP production was implicated in the regulation of macrophage efferocytosis^37^. To assess the contribution of mitochondrial activity, we pretreated BMDMs with oligomycin (1 μM), an ATP synthase inhibitor, or rotenone plus antimycin A (Rot/AA; 0.5 μM each), inhibitors of mitochondrial electron transport, for 30 min. BMDMs were subsequently incubated with fluorescently labeled apoptotic Jurkat cells at a 1:1 ratio for 30 min at 37°C (Fig. 4A).

**Figure 4.**
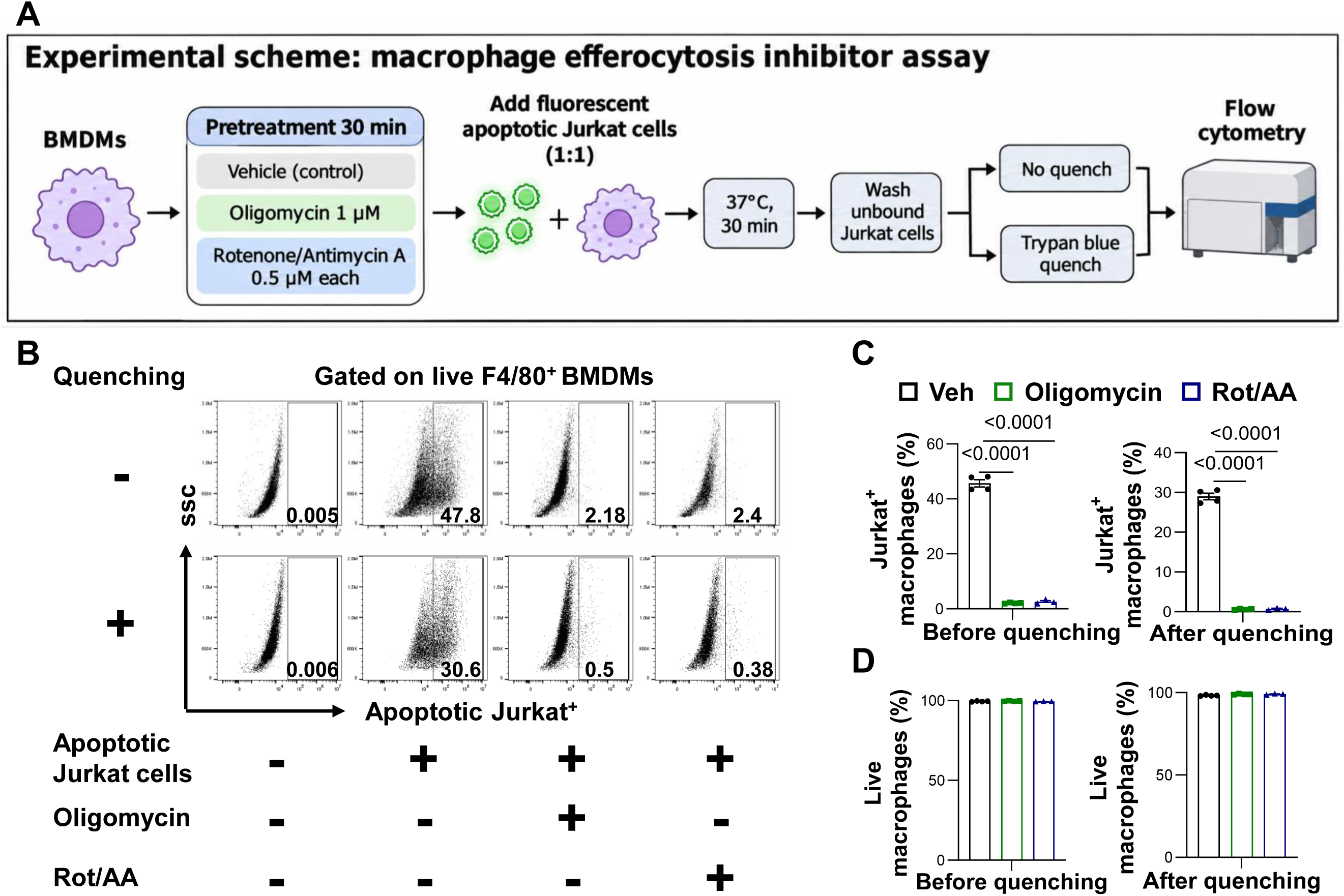
Mitochondrial inhibition markedly suppresses apoptotic-cell-associated fluorescence in BMDMs. Bone marrows were isolated from WT mice and differentiation into BMDMs. (A) Schematic of the apoptotic-cell association assay. BMDMs were pretreated with Veh, oligomycin (1 μM), or Rot/AA (0.5 μM each) for 30 min and incubated with fluorescently labeled apoptotic Jurkat cells at a 1:1 target-to-macrophage ratio for 30 min at 37 °C. Following removal of unbound Jurkat cells, extracellular fluorescence was either left unquenched or quenched with Trypan Blue before flow cytometry. (B) Representative flow cytometry plots showing Jurkat-associated fluorescence in viable F4/80⁺ BMDMs, with and without extracellular quenching. (C) Quantification of Jurkat-positive macrophages before and after quenching in Veh-, oligomycin-, and Rot/AA-treated groups. (D) Percentages of viable macrophages before and after quenching. Data are mean ± SEM (n ≥ 3 per group). Two-tailed unpaired Student’s t-tests for C and D Exact P values are shown where indicated.

Flow cytometric analysis of viable F4/80⁺ macrophages revealed that both oligomycin and Rot/AA markedly reduced apoptotic Jurkat cell-associated fluorescence compared with vehicle-treated controls (Figs. 4B, C). In representative samples, the percentage of Jurkat-positive macrophages decreased from 47.8% in controls to 2.18% and 2.40% following oligomycin and Rot/AA treatment, respectively. To distinguish extracellularly associated fluorescence from internalization-associated signal, we performed extracellular fluorescence quenching using Trypan Blue. Following quenching, the percentage of Jurkat-positive macrophages decreased from 30.6% in controls to 0.50% and 0.38% in the oligomycin-and Rot/AA-treated groups, respectively (Figs. 4B, C). Both inhibitors significantly reduced the percentages of Jurkat-positive macrophages before and after quenching (P < 0.0001).

Importantly, macrophage viability remained comparable among treatment groups (Fig. 4D), suggesting that the reduced apoptotic-cell-associated fluorescence was not attributable to overt cell death. Together, these findings demonstrate that pharmacological inhibition of mitochondrial activity markedly suppresses apoptotic-cell-associated fluorescence, including the quenching-resistant fraction, supporting a role for mitochondrial bioenergetics in macrophage apoptotic-cell uptake.

### 6. Succinate supplementation does not consistently alter macrophage inflammatory gene expression

Given that mDIC regulates mitochondrial dicarboxylate transport, including succinate^38^, we investigated whether altered succinate availability contributes to the macrophage phenotype associated with mDIC deficiency. Extracellular succinate concentrations were comparable between control and mDIC MKO BMDMs (Supplementary Fig. S3A). To further examine the effects of succinate on macrophage activation, BMDMs were treated with succinate (25 μM) for 24 or 48 h, which did not consistently alter the expression of inflammatory or alternative activation-associated genes (Supplementary Fig. S3C). Similarly, pretreatment with succinate at 200 μM or 5 mM did not consistently alter macrophage gene expression following LPS or IL-4 stimulation (Supplementary Fig. S3D–G). Treatment with the cell-permeable succinate derivative diethyl succinate (5 mM) also failed to induce a coordinated transcriptional response under the tested conditions (Supplementary Fig. S3H, I). Together, these findings suggest that extracellular succinate availability and exogenous succinate supplementation do not readily explain the macrophage transcriptional phenotype associated with mDIC deficiency.

### 7. Myeloid mDIC deficiency attenuates diet-induced obesity and promotes an alternative activation-associated macrophage phenotype in adipose tissue

To investigate the physiological consequences of myeloid mDIC deficiency during obesity, control and *mDIC* MKO mice were subjected to high-fat diet (HFD) feeding. Compared with controls, *mDIC* MKO mice exhibited significantly reduced body weight gain, particularly during the later stages of HFD feeding (Fig. 5A). Consistently, insulin tolerance testing revealed lower blood glucose levels in *mDIC* MKO mice following insulin administration, suggesting improved insulin responsiveness (Fig. 5B). Furthermore, *mDIC* MKO mice exhibited significantly reduced SWAT, both in absolute weight (P = 0.003) and relative to body weight (P = 0.0059), whereas epididymal WAT (EWAT) showed a decreasing trend and brown adipose tissue (BAT) mass remained relatively unchanged (Supplementary Fig. S2C).

**Figure 5.**
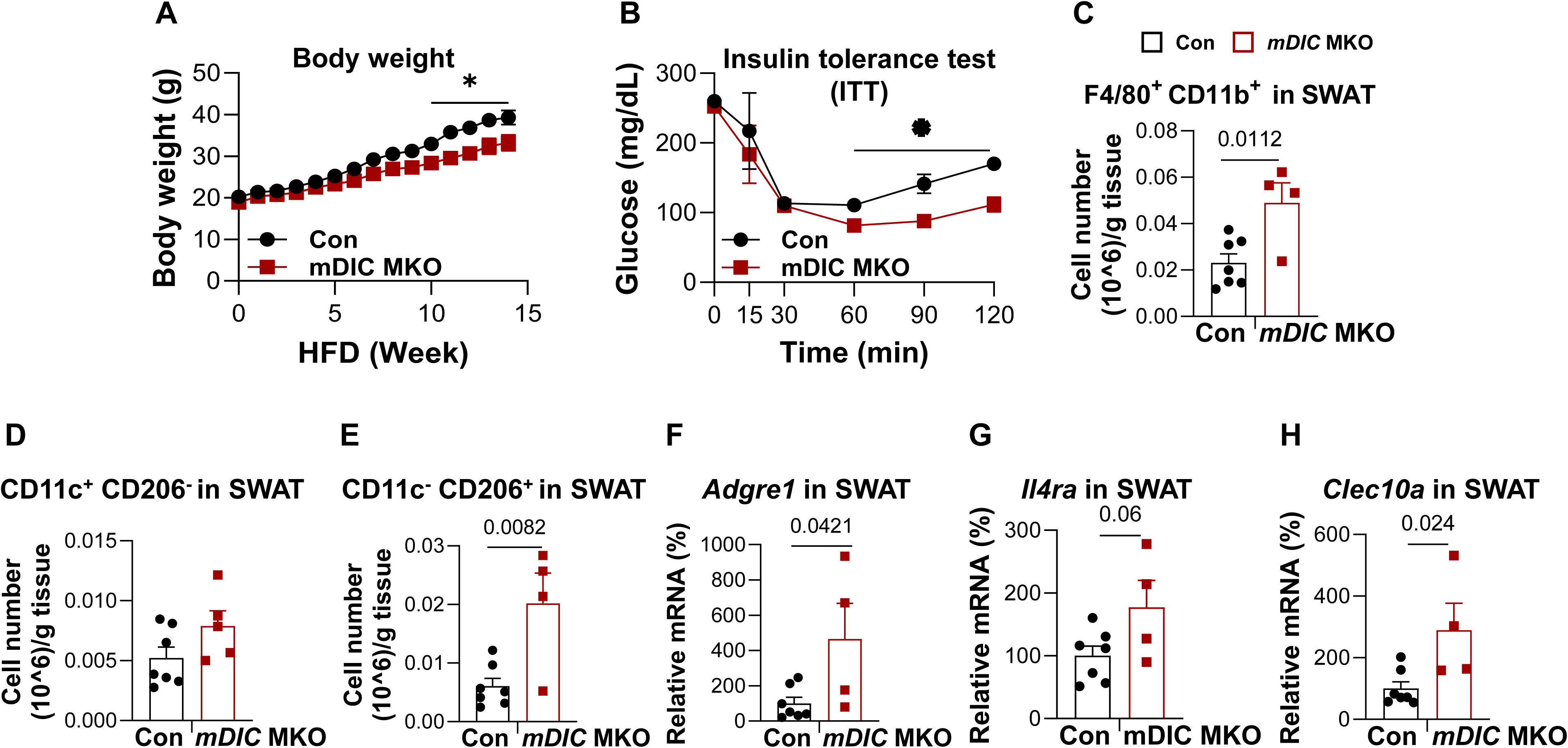
Myeloid mDIC deficiency attenuates diet-induced obesity and remodels adipose macrophage populations. Con and *mDIC* MKO mice were fed a high-fat diet (HFD) for 15 weeks before metabolic and adipose tissue analyses. (A) Body weight trajectories of Con and *mDIC* MKO mice during 15 weeks of HFD feeding (n = 4 per group). (B) Insulin tolerance test (ITT) after 15 weeks of HFD feeding (n = 4 per group). (C–E) Flow cytometric quantification of CD11b⁺F4/80⁺ macrophages (C; n ≥ 4 per group), CD11c⁺CD206⁻ macrophages (D), and CD11c⁻CD206⁺ macrophages (E), expressed as cell number per gram of SWAT. (F–H) Relative mRNA expression of *Adgre1* (F), *Il4ra* (G), and *Clec10a* (H) in SWAT. Data are mean ± SEM. Two-tailed unpaired Student’s t-tests for each time point in A and B and for C–H. Exact P values are shown where indicated; *P < 0.05.

To determine whether these metabolic improvements were accompanied by alterations in adipose macrophage populations, we performed flow cytometric analysis of SWAT. Interestingly, *mDIC* MKO mice exhibited a significantly increased number of CD11b⁺F4/80⁺ macrophages per gram of WAT compared with controls (P = 0.0112; Fig. 5C). Further analysis revealed a significant increase in CD11c⁻CD206⁺ macrophages (P = 0.0082; Fig. 5E), whereas CD11c⁺CD206⁻ macrophages showed a nonsignificant increasing trend (Fig. 5D). These findings suggest that myeloid mDIC deletion alters macrophage abundance and composition in adipose tissue, particularly favoring the accumulation of a CD206-expressing macrophage population.

Consistent with the flow cytometry results, quantitative PCR analysis of SWAT demonstrated significantly increased expression of the macrophage marker *Adgre1* (P = 0.0421) and the alternative activation-associated marker *Clec10a* (P = 0.024) in *mDIC* MKO mice. Expression of *Il4ra* also showed an increasing trend (P = 0.06; Figs. 5F-H). Together, these findings indicate that myeloid mDIC deficiency attenuates HFD-associated metabolic dysfunction and is accompanied by adipose macrophage remodeling characterized by increased CD206⁺ macrophage abundance and alternative activation-associated gene expression.

## Discussion

In the present study, we identified a previously unexplored role for the mitochondrial dicarboxylate carrier (mDIC/SLC25A10) in regulating mitochondrial function in macrophages and adipose immune remodeling during obesity. Our findings demonstrate that palmitate (PA) increases mDIC expression in mouse and human macrophages, whereas myeloid mDIC deletion preserves mitochondrial respiratory capacity under PA stress. Single-cell RNA sequencing further revealed an increased proportion of efferocytosis-associated adipose tissue macrophages (ATMs) in *mDIC* MKO mice, accompanied by changes in macrophage gene expression and apoptotic-cell-associated fluorescence. Pharmacological inhibition of mitochondrial ATP synthase or electron transport markedly reduced apoptotic-cell-associated fluorescence, supporting the involvement of mitochondrial activity in macrophage apoptotic-cell handling. Importantly, myeloid mDIC deletion attenuated high-fat diet (HFD)-induced weight gain, improved insulin tolerance, and increased the abundance of CD11c⁻CD206⁺ macrophages in SWAT. Together, these findings identify myeloid mDIC as a regulator of macrophage metabolic adaptation and adipose immune composition during obesity.

Previous studies have established that mDIC is an important regulator of adipocyte metabolism. An et al. demonstrated that adipocyte mDIC regulates mitochondrial succinate transport and suppresses lipolysis, thereby protecting against hepatic lipotoxicity and systemic insulin resistance^40^. In contrast, the role of mDIC in macrophages has remained largely unexplored. Consistent with previous observations linking reduced adipose mDIC expression to metabolic dysfunction, we observed lower mDIC protein abundance in subcutaneous WAT from individuals with obesity. Interestingly, PA treatment increased mDIC expression in both mouse BMDMs and human THP-1-derived macrophages. These apparently divergent responses may reflect differences in cellular composition, metabolic conditions, or the duration of exposure. Because the human adipose measurements were obtained from whole tissue, they do not establish macrophage-specific changes in mDIC expression during obesity. Nevertheless, these findings suggest that the regulation and physiological consequences of mDIC activity may differ among adipose cell populations. Notably, whereas adipocyte-specific mDIC deletion was previously associated with impaired adipose metabolic function^40^, our myeloid deletion model exhibited a favorable metabolic phenotype during HFD feeding, highlighting the importance of cellular context in determining mDIC function.

One important finding of our study is that mDIC deficiency alters mitochondrial respiration in macrophages under metabolic stress. PA markedly reduced FCCP-stimulated oxygen consumption in control BMDMs, whereas mDIC-deficient macrophages maintained a higher respiratory response. In contrast, basal respiration was broadly comparable across genotypes in the experiment shown. These findings suggest that myeloid mDIC deletion modifies the mitochondrial response to PA rather than constitutively increasing mitochondrial respiration. Consistently, several mitochondrial respiratory chain-associated genes and *Pgc1a* exhibited increased expression trends in mDIC MKO BMDMs. However, these transcriptional changes do not establish increased mitochondrial biogenesis or respiratory complex activity. Moreover, the respiratory phenotype was not uniformly preserved following LPS stimulation, suggesting stimulus-dependent metabolic adaptation. Because mDIC mediates mitochondrial dicarboxylate transport, its deletion may alter mitochondrial substrate availability or compensatory metabolic pathways, although the underlying mechanisms remain to be determined.

Our single-cell transcriptomic analysis provides additional evidence that myeloid mDIC influences adipose macrophage heterogeneity during onset of obesity. After five weeks of HFD feeding, *mDIC* MKO mice exhibited an increased relative abundance of an efferocytosis-associated ATM population, accompanied by reduced proportions of inflammatory ATMs and LAMs. These observations extend previous studies demonstrating that ATMs comprise transcriptionally and functionally heterogeneous populations involved in lipid metabolism, inflammation, and tissue homeostasis^1,11,42–44^. The efferocytosis-associated ATM population exhibited expression of genes associated with tissue homeostasis and lipid handling, including *Mrc1* and *Lpl*. In parallel, mDIC-deficient BMDMs showed increased *Mertk* expression by qPCR and higher expression of several alternative activation-associated genes, particularly *Arg1* and *Mrc1*. Because MerTK is an established receptor involved in apoptotic-cell clearance^18,27^, these observations support the possibility that mDIC deficiency favors a macrophage state associated with efferocytic function. Nevertheless, transcriptional signatures alone cannot establish functional efferocytosis, and further studies are required to determine whether these macrophage populations exhibit enhanced internalization or degradation of apoptotic cells.

Mitochondrial metabolism has increasingly been recognized as an important regulator of macrophage efferocytosis^30,34,35,37^. These findings provide a rationale for investigating whether the mitochondrial respiratory changes associated with mDIC deletion contribute to apoptotic-cell handling. Consistent with an involvement of mitochondrial activity, we observed that oligomycin and Rot/AA markedly suppressed apoptotic Jurkat cell-associated fluorescence in control BMDMs. Importantly, the inhibitory effects persisted following extracellular fluorescence quenching, suggesting a reduction in the internalization-associated fraction rather than only extracellular target binding. Macrophage viability remained comparable among treatment groups, arguing against overt cell death as the primary explanation. However, these observations do not establish that enhanced mitochondrial respiration mediates the macrophage phenotype caused by mDIC deficiency. The inhibitor experiments were conducted in control macrophages, and the near-complete suppression of fluorescence introduces a potential floor effect. Furthermore, mitochondrial metabolism can regulate different stages of efferocytosis, including uptake, phagosome maturation, and anti-inflammatory signaling. Indeed, Morioka et al. demonstrated an efferocytosis-associated glycolytic program with suppression of oxidative phosphorylation^36^, indicating that mitochondrial respiratory requirements may vary with experimental context. Future studies comparing control and mDIC-deficient macrophages under matched mitochondrial inhibition, together with direct measurements of internalization and cargo degradation, will be necessary to establish the metabolic mechanisms underlying the mDIC-associated efferocytic phenotype.

Because mDIC transports dicarboxylates, including succinate and malate, across the inner mitochondrial membrane^38–40^, we also investigated whether succinate contributes to the observed macrophage phenotype. Previous work identified a role for mDIC-dependent succinate export in regulating adipocyte lipolysis^40^. Intracellular succinate has also been implicated in inflammatory macrophage responses through HIF-1α-dependent signaling and succinate dehydrogenase-linked mitochondrial activity^45,46^. Extracellular succinate can also signal through the SUCNR1 receptor^47^. Succinate is also suggested to promote an anti-inflammatory phenotype in adipose tissue macrophages in a succinate receptor 1 (SUCNR1)-dependent manner^48^. However, our *in vitro* results showed that extracellular succinate concentrations were comparable between control and mDIC MKO BMDMs (Supplementary Fig. S3A). Furthermore, supplementation with succinate or diethyl succinate did not consistently induce a coordinated inflammatory or alternative activation-associated transcriptional response under the tested conditions (Supplementary Fig. S3B–I). These findings suggest that changes in extracellular succinate availability alone may not explain the macrophage gene-expression phenotype associated with mDIC deletion. Nevertheless, extracellular succinate concentrations do not directly reflect mitochondrial or cytosolic succinate pools, and exogenous supplementation may not reproduce the metabolic consequences of altered mitochondrial transport. Therefore, a contribution of intracellular succinate metabolism cannot be fully excluded.

Importantly, the macrophage changes associated with myeloid mDIC deletion were accompanied by favorable systemic metabolic outcomes during HFD feeding. *mDIC* MKO mice exhibited reduced body weight gain and lower glucose concentrations during insulin tolerance testing, consistent with improved insulin responsiveness. These changes were accompanied by reduced adipose tissue mass and increased numbers of CD11b⁺F4/80⁺ macrophages per gram of WAT. Further analysis demonstrated increased abundance of CD11c⁻CD206⁺ macrophages, along with higher WAT expression of *Adgre1* and *Clec10a*, and a trend toward increased *Il4ra*. These findings are consistent with remodeling of the adipose macrophage compartment toward an alternative activation-associated phenotype. Prior work has linked macrophage-specific PPARγ signaling to alternative activation and insulin sensitivity during obesity^49^. Interestingly, increased macrophage density occurred despite reduced adipose tissue mass, suggesting that the metabolic benefits of myeloid mDIC deletion are not solely attributable to reduced macrophage accumulation. However, CD206 expression does not uniquely identify efferocytic macrophages, and the current data do not establish that these changes in macrophages directly mediate improved systemic metabolism. Additional studies will be needed to determine the contribution of macrophage efferocytosis and inflammatory resolution to the metabolic phenotype.

Several limitations should be considered. First, LysM-Cre-mediated deletion affects multiple myeloid populations, preventing definitive attribution of the in vivo phenotype to ATMs alone. Second, the single-cell RNA-seq analysis included a limited number of pooled biological samples per genotype, restricting statistical assessment of macrophage subcluster abundance. Third, fluorescence-based apoptotic-cell association assays do not fully distinguish among the steps of efferocytosis, such as surface binding, internalization, and intracellular degradation. Fourth, the mitochondrial inhibitor experiments cannot establish a genotype-specific requirement for respiration in mDIC-regulated efferocytosis. Fifth, although we examined extracellular succinate concentrations and the effects of succinate supplementation, we did not directly assess intracellular succinate metabolism or malate metabolism. Given that mDIC also transports malate^38^, altered malate transport may contribute to the mitochondrial respiratory phenotype observed in mDIC-deficient macrophages. Future metabolomic and metabolic flux analyses will be necessary to clarify the contributions of succinate and malate transport. Finally, differences in body weight and adiposity may contribute to the observed insulin-sensitive phenotype independently of macrophage clearance activity. Future studies using stage-resolved efferocytosis assays, mitochondrial functional measurements, and targeted perturbation of candidate efferocytic regulators, including Neuropilin 2 (NRP2)^50^, will be important for clarifying the relationship between mDIC, macrophage metabolism, and adipose inflammatory resolution.

In conclusion, our study identifies myeloid mDIC as a previously unrecognized regulator of macrophage mitochondrial responses and adipose immune remodeling during obesity. Myeloid mDIC deficiency is associated with preserved mitochondrial respiratory capacity under PA stress, enrichment of efferocytosis-associated ATMs, and improved metabolic outcomes during HFD feeding. These findings suggest a potential connection among mitochondrial dicarboxylate transport, macrophage apoptotic-cell handling, and metabolic homeostasis, providing a foundation for future studies to elucidate the underlying mechanisms and their contributions to obesity-associated cardiometabolic dysfunction.

## Materials and Methods

### Human subcutaneous adipose tissue specimens

De-identified frozen human SWAT specimens were obtained from the Human Metabolic Tissue Bank Core of the University of Pennsylvania Diabetes Research Center (NIH DK019525), shipped on dry ice, and stored at −80 °C until analysis. Samples were from female donors classified as lean or obese, spanning younger and older age groups. The cohort comprised eight lean donors and seven donors with obesity. Available participant characteristics are summarized in Table S1. Upon receipt, samples were stored at −80°C until protein extraction and immunoblot analysis.

### Mouse models and dietary intervention

Myeloid-specific mDIC knockout (mDIC MKO) mice were generated by crossing mice carrying loxP-flanked *Slc25a10* alleles (*Slc25a10*^fl/fl^) with LysM-Cre mice^41^. *Slc25a10*^fl/fl^ littermates lacking Cre served as controls (Con). Deletion of mDIC was verified in bone marrow-derived macrophages (BMDMs) by reverse-transcription quantitative PCR (RT-qPCR) and immunoblotting. C57BL/6J wild-type (WT) mice were purchased from The Jackson Laboratory (Bar Harbor, ME, USA). All animal experiments were approved by the Institutional Animal Care and Use Committee of UTHealth Houston (protocol AWC-24-0103). Mice were maintained on a C57BL/6 genetic background in a specific pathogen-free facility under a 12-h light/12-h dark cycle with ad libitum access to food and water. Male mice were generally enrolled at 8–10 weeks of age. Animals were euthanized by CO₂ inhalation before tissue collection. Primers for genotyping are summarized in Table S2.

Mice received standard chow (LabDiet 5058) or a high-fat diet (HFD; 60% of calories from fat; D12492, Research Diets). Control and *mDIC* MKO mice within each cohort received the same diet. Body weight was recorded weekly. For single-cell RNA sequencing, SWAT was collected after five weeks of HFD feeding. In the long-term obesity study, mice were fed HFD for 15 weeks, underwent insulin tolerance testing, and were subsequently euthanized for tissue collection. SWAT, epididymal WAT (EWAT), and brown adipose tissue (BAT) were weighed; depot masses were normalized to body weight where indicated. SWAT was further analyzed by flow cytometry and RT-qPCR. Male mice were used for all animal experiments unless otherwise specified.

### Insulin tolerance testing

Following a 5-h fast with free access to water, baseline blood glucose was measured from tail-vein blood using a Contour Next glucometer (84413500). Insulin (I5500, MilliporeSigma) was administered by intraperitoneal injection at 1.0 U/kg body weight. Blood glucose concentrations were measured at 15, 30, 45, 60, 90, and 120 min after insulin administration.

### Isolation and differentiation of murine bone marrow-derived macrophages

BMDMs were isolated from mice maintained on either standard chow or a high-fat diet (HFD), as specified for individual experiments. Femurs and tibiae were aseptically collected from euthanized mice after spraying the animals with 70% ethanol. Surrounding tissues were removed and the bone ends opened. Bone marrow was flushed from the shafts using a needle and syringe containing RPMI 1640 medium. Red blood cells were lysed using ACK Lysing Buffer (118-156-101, Quality Biological). The cell suspension was passed through a 100-μm strainer, washed, and centrifuged at approximately 300 × g for 5–10 min.

Bone marrow cells were differentiated into macrophages for seven days in RPMI 1640 (R8758-500ML, MilliporeSigma) supplemented with 10% fetal bovine serum (FBS; F0926-500ML, MilliporeSigma) and 20 ng/mL recombinant mouse macrophage colony-stimulating factor (M-CSF; HZ-1192, Proteintech). Culture medium was replaced every three days^51^. Differentiated BMDMs were harvested and replated for subsequent experiments. For fatty-acid treatment, cells were exposed to palmitic acid (PA; 100 μM for 24 h; HY-N0830, MedChemExpress) or a matched bovine serum albumin (BSA) vehicle control. For mitochondrial respiration experiments involving inflammatory stimulation, BMDMs were treated with lipopolysaccharide (LPS; 10 ng/mL for 24 h; Sigma-Aldrich).

### Human THP-1-derived macrophages

THP-1 monocytes were maintained in RPMI 1640 (R8758-500ML, MilliporeSigma) supplemented with 10% FBS (F0926-500ML, MilliporeSigma). Differentiation was induced with 50 nM phorbol 12-myristate 13-acetate (PMA; HY-18739, MedChemExpress) for 72 h, followed by replacement with PMA-free complete medium and a further 24-h rest. Differentiated THP-1 macrophages were exposed to PA (100 μM for 24 h; HY-N0830, MedChemExpress) or a matched BSA control before analysis of SLC25A10 mRNA and mDIC protein expression.

### RNA extraction and quantitative real-time PCR

Adipose tissues were snap-frozen in liquid nitrogen and stored at −80 °C. Frozen tissues were homogenized using a TissueLyser II (Qiagen). Total RNA from tissues and cultured macrophages was extracted using the RNeasy Mini Kit (74106, Qiagen) according to the manufacturer’s instructions. RNA concentration and purity were assessed using a NanoDrop spectrophotometer (Thermo Fisher Scientific). One microgram of total RNA was reverse-transcribed with the HiScript IV First-Strand cDNA Synthesis Kit with gDNA wiper (R412-02, Vazyme). Quantitative PCR was performed using SYBR Green PCR Master Mix (RK21203, ABclonal Technology) on a CFX Opus 384 real-time PCR system (Bio-Rad). Relative transcript abundance was calculated by the 2^(−ΔΔCt) method; Stx5a was used as the reference gene for mouse samples and *RPL37A* for human THP-1-derived macrophages. Primers for QPCR are summarized in Table S3.

### Protein extraction and immunoblotting

Frozen sWAT and cultured macrophages were lysed in T-PER Tissue Protein Extraction Reagent (78510, Thermo Fisher Scientific) supplemented with protease and phosphatase inhibitors (P3100-005, GenDEPOT). Tissue lysates were clarified by centrifugation at 20,000 × g for 15 min at 4 °C, and protein concentrations were determined using a bicinchoninic acid assay (23225, Thermo Fisher Scientific). Equal amounts of protein were separated on 4–12% SDS–polyacrylamide gels (Bio-Rad) and transferred to nitrocellulose membranes. Membranes were blocked with 5% BSA in Tris-buffered saline containing Tween-20 and incubated overnight at 4 °C with rabbit anti-SLC25A10/mDIC (12086-1-AP, Proteintech; 1:1,000) or mouse anti-β-actin (A3854, MilliporeSigma; 1:5,000). Membranes were washed and incubated for 1 h at room temperature with anti-rabbit IgG-HRP (7074S, Cell Signaling Technology; 1:2,000) or goat anti-mouse IgG-HRP (A16066, Invitrogen; 1:100,000), as appropriate. Chemiluminescent bands were visualized using a ChemiDoc Touch imaging system (Bio-Rad); band intensities were quantified using ImageJ and normalized to β-actin.

### Mitochondrial respiration measurements

Oxygen consumption rate (OCR) was measured in Con and *mDIC* MKO BMDMs using a Seahorse XFe24 Extracellular Flux Analyzer (Agilent Technologies). Before analysis, cells were equilibrated for 45–60 min at 37 °C in a non-CO₂ incubator in Seahorse XF assay medium (103575-100, Agilent Technologies) containing 10 mM glucose (103577-100), 2 mM glutamine (103579-100), and 1 mM sodium pyruvate (103578-100). Basal OCR was recorded, followed by sequential injections of oligomycin (2.5 μM), FCCP (2 μM), and rotenone plus antimycin A (1.0 μM each), using the Seahorse XF Cell Mito Stress Test Kit (103016-100, Agilent Technologies)^52^. Concentrations refer to the final assay-well concentrations. OCR was normalized to total protein per well. Cells were assessed following PA (100 μM for 24 h), LPS (10 ng/mL for 24 h), or the corresponding control treatment.

### Adipose stromal vascular fraction isolation and single-cell RNA sequencing

For single-cell RNA sequencing (scRNA-seq), SWAT was collected after five weeks of HFD feeding, rinsed in phosphate-buffered saline (PBS), and minced into approximately 1-mm² fragments. Tissue was digested for 30–45 min at 37 °C with agitation at 200 rpm in PBS containing 10 mM CaCl₂, collagenase D (1.5 U/mL; 11088882001, MilliporeSigma), and Dispase II (2.4 U/mL; 04942078001, MilliporeSigma). Digested tissue was filtered through a 100-μm strainer (76327-102, VWR), centrifuged at 300 × g for 5 min, and resuspended. Residual erythrocytes were removed with ACK Lysing Buffer (118-156-101, Quality Biological). The resulting suspension was passed through a 40-μm strainer (76327-098, VWR) and centrifuged at 300 × g for 5 min. Stromal vascular fraction (SVF) cells were resuspended as a single-cell suspension for sequencing.

Single-cell transcriptomes were profiled using Parse Biosciences Evercode Whole Transcriptome technology. Cells were fixed and permeabilized for split-pool combinatorial barcoding, followed by library preparation and next-generation sequencing. Sequencing reads were processed to generate cell-by-gene count matrices. Downstream analyses were performed in R. Following quality filtering, data were normalized, dimensionally reduced, and clustered, and uniform manifold approximation and projection (UMAP) was used for visualization. Major cell populations were annotated using established marker genes. The monocyte/macrophage compartment was isolated and re-clustered to characterize inflammatory adipose tissue macrophages (ATMs), lipid-associated macrophages (LAMs), efferocytosis-associated ATMs, transitional LAMs, proliferating ATMs, and infiltrating monocytes. Cell proportions were summarized by pooled biological sample.

### Flow cytometric characterization of adipose tissue macrophages

SVF cell suspensions from SWAT were incubated with Fc receptor-blocking anti-mouse CD16/32 (clone 93, 101302, BioLegend; 1:400) and stained with PE/Cyanine7-conjugated anti-mouse CD45 (clone 30-F11, BioLegend), Pacific Blue-conjugated anti-mouse/human CD11b (clone M1/70, 101224, BioLegend; 1:100), PE-conjugated anti-mouse F4/80 (clone BM8, 123110, BioLegend; 1:100), FITC-conjugated anti-mouse CD206 (clone C068C2, 141704, BioLegend; 1:100), and APC-conjugated anti-mouse CD11c (clone N418, 117310, BioLegend; 1:100). Following exclusion of debris and doublets, viable CD45-positive leukocytes were identified, followed by gating of CD11b-positive F4/80-positive macrophages. CD11c-positive CD206-negative and CD11c-negative CD206-positive subpopulations were quantified within the macrophage gate. Samples were acquired using a CytoFLEX LX flow cytometer (Beckman Coulter), and macrophage numbers were expressed per gram of SWAT.

### Generation, apoptosis assessment, and fluorescent labeling of Jurkat cells

Jurkat T cells were maintained in RPMI 1640 (R8758-500ML, MilliporeSigma) containing 10% FBS (F0926-500ML, MilliporeSigma). Viability was assessed before irradiation, and only cultures with at least 95% viable cells were used. Apoptosis was induced by 10 Gy X-ray irradiation followed by a 16-h recovery in complete medium. Apoptosis was assessed in a separate aliquot by Annexin V-FITC and propidium iodide (PI) staining according to the manufacturer’s protocol (Sigma-Aldrich). Annexin V-positive/PI-negative and Annexin V-positive/PI-positive populations were used to identify early apoptotic and late apoptotic or membrane-compromised cells, respectively.

For flow-cytometric apoptotic-cell association assays, Jurkat cells were labeled with PKH67 Green Fluorescent Cell Linker Midi Kit (MIDI67, Sigma-Aldrich); BMDMs were identified separately by anti-F4/80 staining. For microscopy assays, BMDMs were labeled with green PKH67 and apoptotic Jurkat cells with red PKH26 (MIDI26, Sigma-Aldrich). For membrane dye labeling, a working dye mixture was prepared using 0.4 μL dye per 100 μL Diluent C. Cells were labeled using the manufacturer’s membrane-labeling workflow, including removal of serum before labeling, reaction termination with serum or complete medium, and washing to remove unbound dye.

### Fluorescence microscopy of macrophage–apoptotic-cell association

PKH67-labeled Con and *mDIC* MKO BMDMs were incubated with PKH26-labeled apoptotic Jurkat cells at a 1:1 target-to-macrophage ratio for 30 min at 37 °C. Cultures were washed with PBS to remove unbound Jurkat cells and imaged using a Nikon AX R confocal microscope. Overlapping red and green fluorescence was quantified using ImageJ relative to macrophage-associated green fluorescence. Eight BMDM samples per genotype were examined, with at least three microscopic fields obtained per sample; additional quantification was performed in three independently prepared BMDM samples. Multiple fields from a single sample were treated as nested observations rather than independent biological replicates.

### Mitochondrial inhibitor treatment and efferocytosis-associated flow cytometry

WT BMDMs were pretreated for 30 min with vehicle, oligomycin (1 μM), or rotenone plus antimycin A (Rot/AA; 0.5 μM each). Inhibitor-containing medium was removed, and PKH67-labeled apoptotic Jurkat cells were added at a 1:1 target-to-macrophage ratio for 30 min at 37 °C. Cultures were washed with PBS to remove unbound Jurkat cells. Macrophages were subsequently stained with eBioscience Fixable Viability Dye eFluor 450 (65-0863-18, Invitrogen/Thermo Fisher Scientific; 1:1,000) and PE-conjugated anti-mouse F4/80 (clone BM8, 123110, BioLegend; 1:100). Flow cytometry was performed using a CytoFLEX LX flow cytometer (Beckman Coulter); singlets and viable F4/80-positive macrophages were gated to determine the proportion positive for PKH67-labeled apoptotic Jurkat cells.

To assess the extracellular contribution to Jurkat-associated fluorescence, parallel samples were analyzed with and without extracellular fluorescence quenching using 0.04% Trypan Blue. Unquenched PKH67 fluorescence was used to assess total apoptotic-cell-associated signal, whereas quenching-resistant fluorescence was used as a surrogate for internalization-associated signal. Macrophage viability was assessed in parallel using eFluor 450.

### Extracellular succinate measurement

Conditioned medium from Con and *mDIC* MKO BMDMs was collected for measurement of extracellular succinate using the EnzyChrom Succinate Assay Kit (ESNT-100, BioAssay Systems), following the manufacturer’s microplate procedure. Briefly, 20 μL of medium sample was combined with 80 μL of freshly prepared working reagent and incubated for 30 min at room temperature. Succinate concentrations were calculated using the kit’s calibration procedure and adjusted for sample dilution. Values were expressed as μM and represent extracellular succinate rather than intracellular metabolite content.

### Succinate and diethyl succinate supplementation

For supplementation experiments, BMDMs were treated with succinate (25 μM) for 24 or 48 h without subsequent stimulation. In separate experiments, cells were pretreated with succinate (200 μM or 5 mM) for 3 h before stimulation with LPS (100 ng/mL) or interleukin-4 (IL-4; 20 ng/mL) for 12 h. Additional cultures were pretreated with diethyl succinate (5 mM) for 90 min, followed by LPS (100 ng/mL) or IL-4 (20 ng/mL) stimulation for 24 h. Expression of inflammatory and alternative activation-associated genes was analyzed by RT-qPCR.

### Statistical analysis

Data are presented as mean ± standard error of the mean (SEM). Statistical analyses were performed using GraphPad Prism (GraphPad Software). Two independent groups were compared using two-sided unpaired Student’s t-tests where applicable. Experiments involving two independent factors and OCR time courses were assessed using two-way analysis of variance (ANOVA), as appropriate. Each mouse or independently prepared macrophage sample was considered a biological replicate. The two pooled single-cell libraries per genotype were treated as independent samples; multiple microscopic fields from one culture were not considered independent biological replicates. Statistical significance was defined as P < 0.05.

## Acknowledgments

This work was supported by National Institutes of Health (NIH) grants K01-DK125447, R03-DK135783, and P30-DK056338 (Pilot Award) to Y.A.A. and American Diabetes Association (ADA) Postdoctoral Fellowship Award 11-23-PDF-23 to F.W. We have used ChatGPT and BioRender to generate schematic figures.

## Author Contributions

F.W. and Y.A.A. conceived the study. F.W. designed and performed experiments, collected and analyzed data, prepared figures, and wrote the original manuscript. P.M.H. and A.S. assisted with experiments and data collection. Q.H. and J.B. provided efferocytosis protocols and reagents. Y.Y. provided training and technical guidance in flow cytometry. X.M. contributed to data interpretation and critically reviewed and edited the manuscript. J.B. provided scientific guidance and resources related to efferocytosis. Y.A.A. supervised the study, provided scientific guidance and resources, contributed to data interpretation, and reviewed and edited the manuscript.

## Figure Legends

**Supplementary Figure S1. mDIC expression in human adipose tissue and mitochondrial respiration following LPS stimulation.**

(A) Representative mDIC and β-actin immunoblots of human SWAT from lean donors (n = 8) and donors with obesity (n = 7), with densitometric quantification (P = 0.0263). (B) Seahorse OCR profiles of Con and *mDIC* MKO BMDMs with or without lipopolysaccharide (LPS; 10 ng/mL, 24 h; n ≥ 4 per group). Oligomycin, FCCP, and Rot/AA were sequentially injected; OCR was normalized to protein content. Data are mean ± SEM. Two-tailed unpaired Student’s t-test for A; two-way ANOVA for B. Exact P values are shown.

**Supplementary Figure S2. Marker gene expression in adipose SVF and macrophage subclusters and adipose tissue weights.**

(A) Violin plots showing expression of representative lineage-marker genes across the major SVF cell populations identified by scRNA-seq: B cells, T cells, CD4⁺ T cells, fibroblasts, macrophages, cycling cells, endothelial cells, monocytes, and mural cells.

(B) Violin plots showing marker genes across six monocyte/macrophage subclusters: inflammatory ATMs, LAMs, efferocytosis-associated ATMs, transitional LAMs, proliferating ATMs, and infiltrating monocytes. (C) Absolute weights of SWAT, epididymal white adipose tissue (EWAT), and brown adipose tissue (BAT) and the corresponding tissue weights normalized to body weight in HFD-fed Con and *mDIC* MKO mice (n = 3 per group). Data are mean ± SEM for C. Two-tailed unpaired Student’s t-tests for C. Exact P values are shown.

**Supplementary Figure S3. Extracellular succinate concentrations and effects of succinate supplementation on macrophage gene expression.**

(A) Extracellular succinate concentrations in conditioned medium from Con and *mDIC* MKO BMDMs (n ≥ 14 per group). (B) Experimental schematic of succinate and diethyl succinate treatments followed by RT-qPCR. (C) Relative mRNA expression of *Arg1*, *Il4*, *Il4ra*, *Il10*, *Mgl2*, *Fizz1*, *Chil3*, *Mrc1*, *Tnfa*, *Il1b*, *iNos*, *Ifn*g, and *Il6* in unstimulated BMDMs treated with succinate (25 μM) for 24 or 48 h (n ≥ 3). (D, E) Expression of *Tnfa*, *Il1b*, *iNos*, *Il6*, and *Ifng* following LPS stimulation (D) or *Arg1*, *Mrc1*, *Il4*, *Il4ra*, *Il10*, *Mgl2*, *Fizz1*, and *Chil3* following IL-4 stimulation (E) in BMDMs pretreated with succinate (200 μM) for 3 h, followed by LPS (100 ng/mL) or IL-4 (20 ng/mL) for 12 h (n = 6). (F, G) Corresponding expression of *Tnfa*, *Il1b*, *iNos*, *Il6*, and *Ifng* in LPS-stimulated BMDMs (F) and *Arg1*, *Mrc1*, *Il4*, *Il4ra*, *Il10*, *Mgl2*, *Fizz1*, and *Chil3* in IL-4-stimulated BMDMs (G) following succinate pretreatment (5 mM, 3 h) and subsequent 12-h stimulation (n = 4). (H, I) Corresponding expression of *Tnfa*, *Il1b*, *iNos*, *Ifng*, and *Il6* in LPS-stimulated BMDMs (H) and *Arg1*, *Mrc1*, *Il4*, *Il4ra*, *Il10*, *Mgl2*, *Fizz1*, and *Chil3* in IL-4-stimulated BMDMs (I) after pretreatment with diethyl succinate (5 mM, 90 min), followed by 24-h stimulation with LPS or IL-4 (n = 4). Data are mean ± SEM. Two-tailed unpaired Student’s t-tests for A and C–I; Exact P values are shown where indicated.

