## Supplementary material for "Myeloid mDIC Deficiency Remodels Adipose Macrophages to an Efferocytosis-Associated State and Improves Metabolic Homeostasis During Obesity": Table S

**Supplementary Materials**

**Tables S1-S3**

**Figures S1-S3**

**Table S1. Characteristics of human SWAT donors**

| Characteristic | Lean (n = 8) | Obesity (n = 7) |
|---|---|---|
| Age, years | 46.5 ± 17.2 (22–70) | 42.0 ± 18.1 (20–66) |
| Young, n (%) | 4 (50.0%) | 4 (57.1%) |
| Aged, n (%) | 4 (50.0%) | 3 (42.9%) |
| Female, n (%) | 8 (100%) | 7 (100%) |
| BMI, kg/m² | 22.71 ± 1.22 (20.76–24.88) | 38.58 ± 1.14 (36.56–39.56) |
| White, n (%) | 6 (75.0%) | 4 (57.1%) |
| Black, n (%) | 1 (12.5%) | 3 (42.9%) |
| Asian, n (%) | 1 (12.5%) | 0 |
| Not Hispanic/Latino, n (%) | 6 (75.0%) | 7 (100%) |
| Hispanic/Latino, n (%) | 1 (12.5%) | 0 |
| Ethnicity not reported, n (%) | 1 (12.5%) | 0 |

**Table S2. Genotyping primers:**

| **Gene** | **Primer** | **Sequence (5′–3′)** |
|---|---|---|
| *mDIC fl/fl* | Forward | GGAGTAGCATATACAGGTCC |
| | Reverse | ACGTCACAGTCTGTCAACAC |
| LysM-Cre | Forward | CACATCTCATGCTGCTCAGATAGCC |
| | Mutant reverse | ACCGGTAATGCAGGCAAAT |
| | Wild-type reverse | GTCACTCACTGCTCCCCTCT |

**Table S3. qPCR primers:**

| Primers | Forward sequence | Reverse sequence |
|---|---|---|
| *Pgc1a* | TCCTCACACCAAACCCACAGAA | TTGGCTTGAGCATGTTGCGA |
| *Cox8a* | TTCCTGCTTCGTGTGTTGTC | GATTGCAGAAGAGGTGACTGG |

| | | |
|---|---|---|
| *Cycs* | CCAGGTATACAAGCAGGTGTGCTC | CATCATTAGGGCCATCCTGGAC |
| *Cox1* | GCACTGGTGGATGCCTTCT | TCTCTCGGGACTCCTTGATGA |
| *Cox2* | AGTTGATAACCGAGTCGTTCTGCCA | TCGGCCTGGGATGGCATCAGT |
| *Cox3* | ACCTACCAAGGCCACCACACTCC | GCAGCCTCCTAGATCATGTGTTGGT |
| *Nd1* | CAGCCGGCCCATTCGCGTTA | AGCGGAAGCGTGGATAGGATGC |
| *Nd2* | TCCTCCTGGCCATCGTACTCAACT | AGAAGTGGAATGGGGCGAGGC |
| *Nd3* | ACCCTACAAGCTCTGCACGCC | GCTCATGGTAGTGGAAGTAGAAGGGCA |
| *Ndufs3* | TGGCAGCACGTAAGAAGGG | CTTCGGTAAGATTTCAGCCACAT |
| *Ndufv1* | CTTCCCCACTGGCCTCAAG | CCAAAACCCAGTGATCCAGC |
| *Stx5a* | CTGAAACAGCAGAGGAACCGTC | GGTCCATCATGTCAATAGCCACG |
| *Slc25a10* | CGAATGACTGGATTGGCACTGC | TGGTCTCGTAGATTGCGAACCG |
| *Mertk* | ATCATCCTCGGCTGCTTCTGTG | ACGACCAGTTGGGAATCCTCCT |
| *Arg1* | CATTGGCTTGCGAGACGTAGAC | GCTGAAGGTCTCTTCCATCACC |
| *Mrc1* | GTTCACCTGGAGTGATGGTTCTC | AGGACATGCCAGGGTCACCTTT |
| *Il4* | *ATCATCGGCATTTTGAACGAGGTC* | *ACCTTGGAAGCCCTACAGACGA* |
| *Il10* | CGGGAAGACAATAACTGCACCC | CGGTTAGCAGTATGTTGTCCAGC |

| | | |
|---|---|---|
| *Clec10a* | ACCCAAGAGCCTGCTAAAGC | GATCCAATCACGGAGCAGC |
| *Mgl2* | CGAGACTTGAGCCAGAAGGTGA | GCCTTCAAGTCTGTCTCCAGCT |
| *Retnla* | CAAGGAACTTCTTGCCAATCCAG | CCAAGATCCACAGGCAAAGCCA |
| *Nos2* | GAGACAGGGAAGTCTGAAGCAC | CCAGCAGTAGTTGCTCCTCTTC |
| *Tnf* | GGTGCCTATGTCTCAGCCTCTT | GCCATAGAACTGATGAGAGGGAG |
| *Il1b* | TGGACCTTCCAGGATGAGGACA | GTTCATCTCGGAGCCTGTAGTG |
| *Il6* | TACCACTTCACAAGTCGGAGGC | CTGCAAGTGCATCATCGTTGTTC |
| *Adgre1* | CGTGTTGTTGGTGGCACTGTGA | CCACATCAGTGTTCCAGGAGAC |
| *Il4ra* | ACCAGATGGAACTGTGGGCTGA | AGCAGCCATTCGTCGGACACAT |
| *Chil3* | TACTCACTTCCACAGGAGCAGG | CTCCAGTGTAGCCATCCTTAGG |
| *Ifng* | CAGCAACAGCAAGGCGAAAAAGG | TTTCCGCTTCCTGAGGCTGGAT |
| *SLC25A10 (human)* | GCAGACTTGGTCAACGTCAGGA | CATGGTTGCACCCGAGAACAGT |
| *Rpl37a (human)* | AATCAGCCAGCACGCCAAGTAC | GCCACTGTCTTCATGCAGGAAC |

### Figure S1

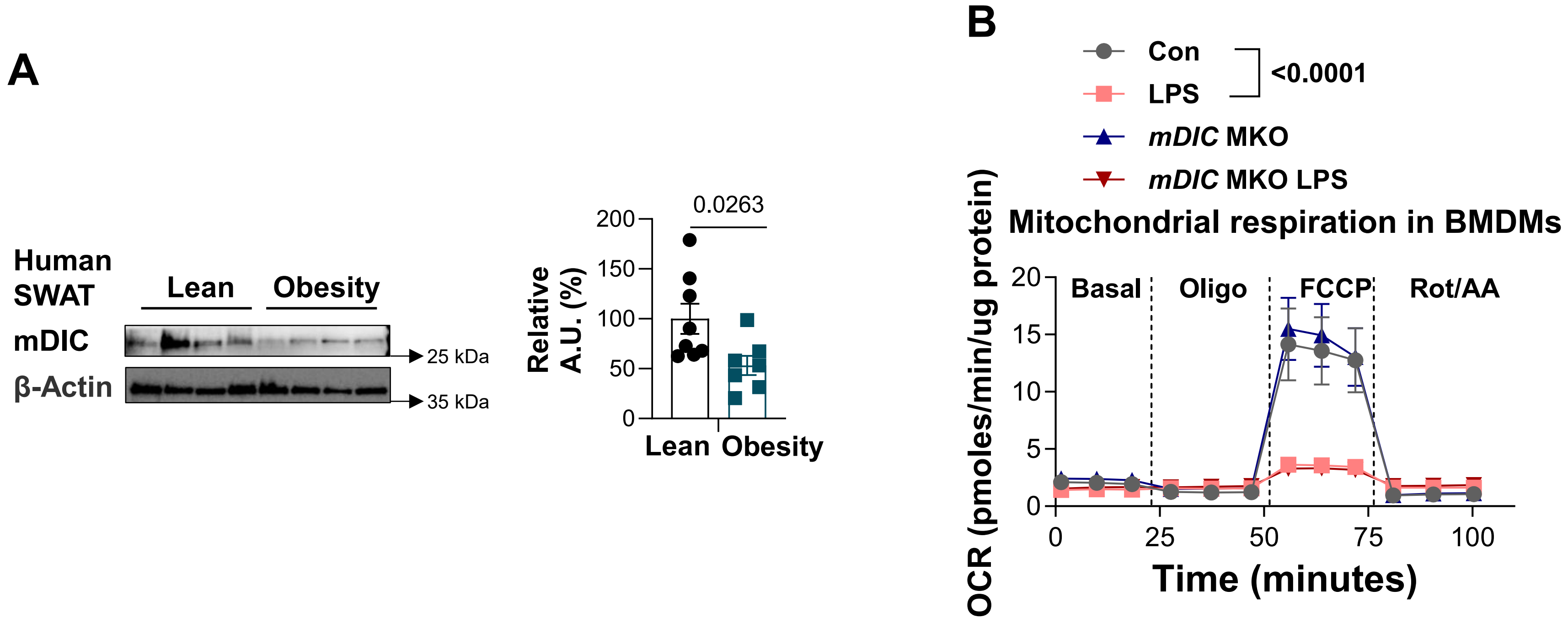


**Supplementary Figure S1. mDIC expression in human adipose tissue and mitochondrial respiration following LPS stimulation.**

(A) Representative mDIC and β-actin immunoblots of human SWAT from lean donors (n = 8) and donors with obesity (n = 7), with densitometric quantification (P = 0.0263). (B) Seahorse OCR profiles of Con and *mDIC* MKO BMDMs with or without lipopolysaccharide (LPS; 10 ng/mL, 24 h; n ≥ 4 per group). Oligomycin, FCCP, and Rot/AA were sequentially injected; OCR was normalized to protein content. Data are mean ± SEM. Two-tailed unpaired Student's t-test for A; two-way ANOVA for B. Exact P values are shown.

Figure S2

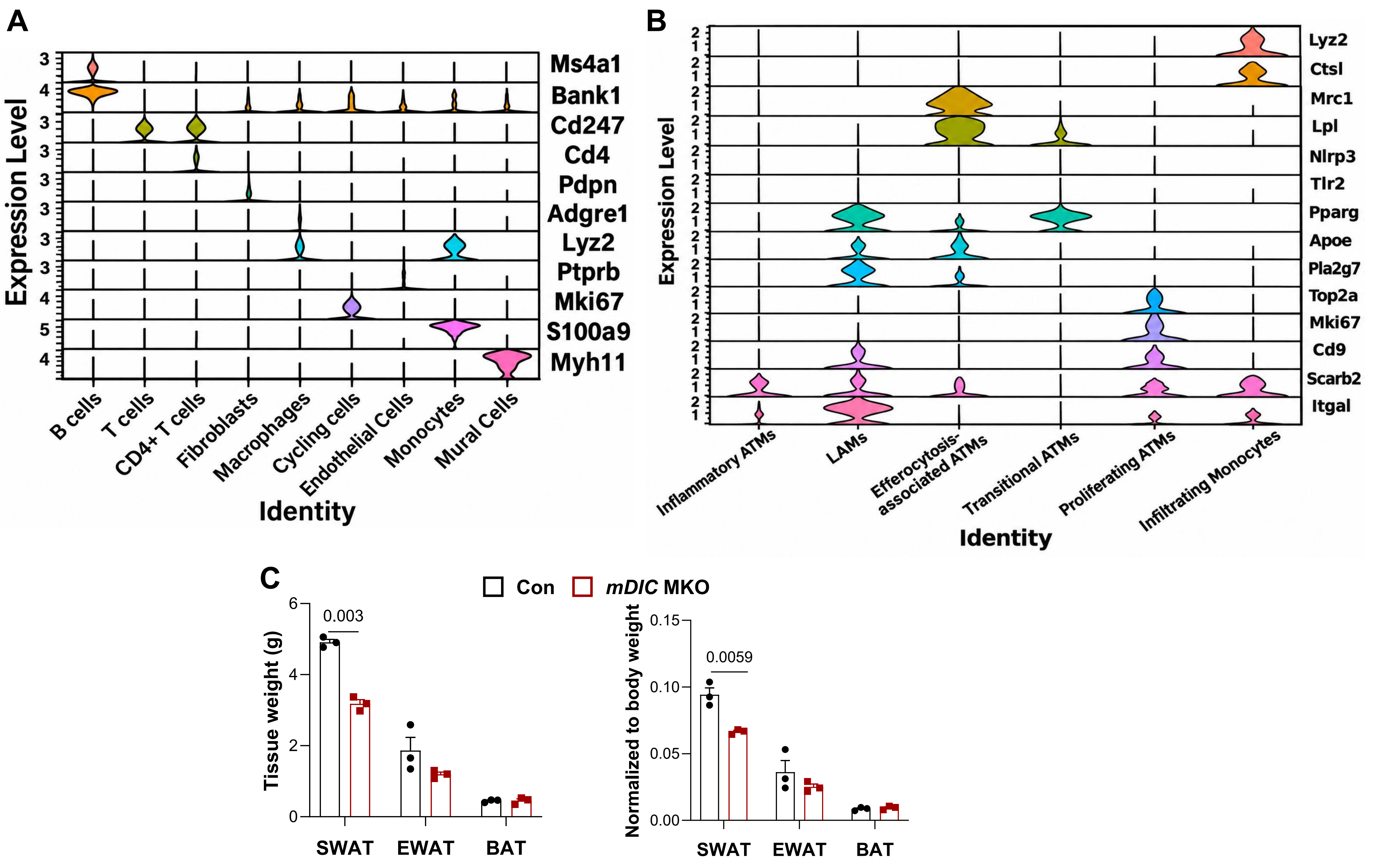
A
Expression Level
Ms4a1
Bank1
Cd247
Cd4
Pdpn
Adgre1
Lyz2
Ptprb
Mki67
S100a9
Myh11
B cells
T cells
CD4+ T cells
Fibroblasts
Macrophages
Cycling cells
Endothelial Cells
Monocytes
Mural Cells
Identity
B
Expression Level
Lyz2
Ctsl
Mrc1
Lpl
Nlrp3
Tlr2
Pparg
Apoe
Pla2g7
Top2a
Mki67
Cd9
Scarb2
Itgal
Inflammatory ATMs
LAMs
Efferocytosis-associated ATMs
Transitional ATMs
Proliferating ATMs
Infiltrating Monocytes
Identity
C
Con
mDIC MKO
Tissue weight (g)
0.003
SWAT
EWAT
BAT
Normalized to body weight
0.0059
SWAT
EWAT
BAT

**Supplementary Figure S2. Marker gene expression in adipose SVF and macrophage subclusters and adipose tissue weights.**
(A) Violin plots showing expression of representative lineage-marker genes across the major SVF cell populations identified by scRNA-seq: B cells, T cells, $CD4^+$ T cells, fibroblasts, macrophages, cycling cells, endothelial cells, monocytes, and mural cells. (B) Violin plots showing marker genes across six monocyte/macrophage subclusters: inflammatory ATMs, LAMs, efferocytosis-associated ATMs, transitional LAMs, proliferating ATMs, and infiltrating monocytes. (C) Absolute weights of SWAT, epididymal white adipose tissue (EWAT), and brown adipose tissue (BAT) and the corresponding tissue weights normalized to body weight in HFD-fed Con and *mDIC* MKO mice (n = 3 per group). Data are mean ± SEM for C. Two-tailed unpaired Student's t-tests for C. Exact P values are shown.

### Figure S3

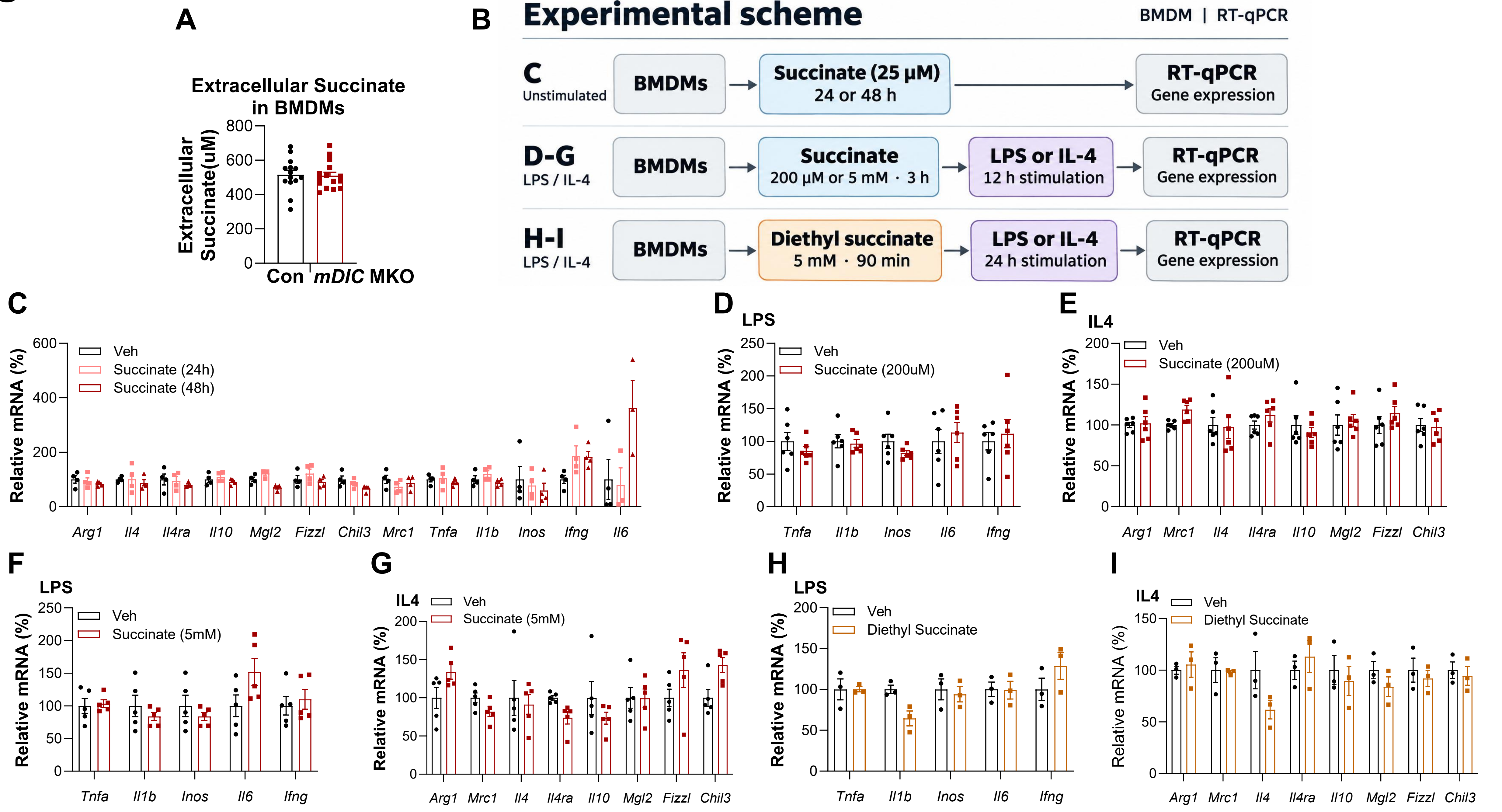
A
Extracellular Succinate in BMDMs
Extracellular Succinate(uM)
800
600
400
200
0
Con
mDIC MKO
B
Experimental scheme
BMDM | RT-qPCR
C
Unstimulated
BMDMs
Succinate (25 μM) 24 or 48 h
RT-qPCR Gene expression
D-G
LPS / IL-4
BMDMs
Succinate 200 μM or 5 mM · 3 h
LPS or IL-4 12 h stimulation
RT-qPCR Gene expression
H-I
LPS / IL-4
BMDMs
Diethyl succinate 5 mM · 90 min
LPS or IL-4 24 h stimulation
RT-qPCR Gene expression
C
Relative mRNA (%)
Veh
Succinate (24h)
Succinate (48h)
Arg1 Il4 Il4ra Il10 Mgl2 Fizzl Chil3 Mrc1 Tnfa Il1b Inos Ifng Il6
D
LPS
Veh
Succinate (200uM)
Tnfa Il1b Inos Il6 Ifng
E
IL4
Veh
Succinate (200uM)
Arg1 Mrc1 Il4 Il4ra Il10 Mgl2 Fizzl Chil3
F
LPS
Veh
Succinate (5mM)
Tnfa Il1b Inos Il6 Ifng
G
IL4
Veh
Succinate (5mM)
Arg1 Mrc1 Il4 Il4ra Il10 Mgl2 Fizzl Chil3
H
LPS
Veh
Diethyl Succinate
Tnfa Il1b Inos Il6 Ifng
I
IL4
Veh
Diethyl Succinate
Arg1 Mrc1 Il4 Il4ra Il10 Mgl2 Fizzl Chil3

**Supplementary Figure S3. Extracellular succinate concentrations and effects of succinate supplementation on macrophage gene expression.**
(A) Extracellular succinate concentrations in conditioned medium from Con and *mDIC* MKO BMDMs (n ≥ 14 per group). (B) Experimental schematic of succinate and diethyl succinate treatments followed by RT-qPCR. (C) Relative mRNA expression of *Arg1*, *Il4*, *Il4ra*, *Il10*, *Mgl2*, *Fizz1*, *Chil3*, *Mrc1*, *Tnfa*, *Il1b*, *iNos*, *Ifn*g, and *Il6* in unstimulated BMDMs treated with succinate (25 μM) for 24 or 48 h (n ≥ 3). (D, E) Expression of *Tnfa*, *Il1b*, *iNos*, *Il6*, and *Ifng* following LPS stimulation (D) or *Arg1*, *Mrc1*, *Il4*, *Il4ra*, *Il10*, *Mgl2*, *Fizz1*, and *Chil3* following IL-4 stimulation (E) in BMDMs pretreated with succinate (200 μM) for 3 h, followed by LPS (100 ng/mL) or IL-4 (20 ng/mL) for 12 h (n = 6). (F, G) Corresponding expression of *Tnfa*, *Il1b*, *iNos*, *Il6*, and *Ifng* in LPS-stimulated BMDMs (F) and *Arg1*, *Mrc1*, *Il4*, *Il4ra*, *Il10*, *Mgl2*, *Fizz1*, and *Chil3* in IL-4-stimulated BMDMs (G) following succinate pretreatment (5 mM, 3 h) and subsequent 12-h stimulation (n = 4). (H, I) Corresponding expression of *Tnfa*, *Il1b*, *iNos*, *Ifng*, and *Il6* in LPS-stimulated BMDMs (H) and *Arg1*, *Mrc1*, *Il4*, *Il4ra*, *Il10*, *Mgl2*, *Fizz1*, and *Chil3* in IL-4-stimulated BMDMs (I) after pretreatment with diethyl succinate (5 mM, 90 min), followed by 24-h stimulation with LPS or IL-4 (n = 4). Data are mean ± SEM. Two-tailed unpaired Student's t-tests for A and C–I; Exact P values are shown where indicated.
